# Modeling of hepatitis B virus infection spread in primary human hepatocytes

**DOI:** 10.1101/2025.02.05.636596

**Authors:** Zhenzhen Shi, Masataka Tsuge, Nicholson Collier, Yasue Takeuchi, Takuro Uchida, Carolyn M. Rutter, Yuji Teraoka, Susan Uprichard, Yuji Ishida, Chise Tateno, Jonathan Ozik, Harel Dahari, Kazuaki Chayama

## Abstract

Chronic hepatitis B virus (HBV) infection poses a significant global health threat, causing severe liver diseases including cirrhosis and hepatocellular carcinoma. We characterized HBV DNA kinetics in primary human hepatocytes (PHH) over 32 days post-inoculation (pi) and used agent-based modeling (ABM) to gain insights into HBV lifecycle and spread. Parallel PHH cultures were mock-treated or HBV entry inhibitor Myr-preS1 (6.25 μg/mL) was initiated 24h pi. In untreated PHH, 3 viral DNA kinetic patterns were identified: (1) an initial decline, followed by (2) rapid amplification, and (3) slower amplification/accumulation. In the presence of Myr-preS1, viral DNA and infected cell numbers in phase 3 were effectively blocked, with minimal to no increase. This suggests that phase 2 represents viral amplification in initially infected cells, while phase 3 corresponds to viral spread to naïve cells. The ABM reproduced well the HBV kinetic patterns observed and predicted that the viral eclipse phase lasts between 18 and 38 hours. After the eclipse phase, the viral production rate increases over time, starting with a slow production cycle of 1 virion per day, which gradually accelerates to 1 virion per hour after 3 days. Approximately 4 days later, virion production reaches a steady state production rate of 4 virions/hour. The estimated median efficacy of Myr-preS1 in blocking HBV spread was 91% (range: 90-92%). The HBV kinetics and the predicted estimates of the HBV eclipse phase duration and HBV production cycles in PHH are similar of those predicted in uPA/SCID mice with human livers.

**Importance:** While primary human hepatocytes (PHH) are the most physiologically relevant culture system for studying HBV infection *in vitro*, comprehensive understanding of HBV infection kinetics and spread in PHH is lacking. In this study, we characterize HBV viral kinetics and employ agent-based modeling (ABM) to provide quantitative insights into the HBV production cycle and viral spread in PHH. The ABM provides an estimate of HBV eclipse phase duration, HBV production cycles and Myr-preS1 efficacy in blocking HBV spread in PHH. The results resemble those predicted in uPA/SCID mice with human livers, demonstrating that estimated HBV infection kinetic parameters in PHH *in vitro* mirrors that observed in *in vivo* HBV infection chimeric mouse model.

## Introduction

An estimated 254 million people are chronically infected with hepatitis B virus (HBV), with 1.2 million new infections reported annually (1). HBV infection leads to approximately 1.1 million deaths each year, primarily due to HBV-related liver disease such as cirrhosis and hepatocellular carcinoma (1). Current treatment options for chronic hepatitis B include antiviral therapy with nucleoside/nucleotide analogues (NUCs) and/or pegylated interferon alpha (2, 3). Although these treatments can suppress HBV replication and slow the progression of liver disease, they do not cure the infection. To develop more effective antiviral therapies aimed at eradicating HBV, it is essential to better understand HBV lifecycle and spread (4).

Primary human hepatocytes (PHH) are well known for their high susceptibility to HBV infection and their utility in molecular studies of HBV lifecycle (5). However, only a few studies have utilized PHH cultures to examine HBV viral kinetics, and comprehensive data on viral dynamics and the kinetics of HBV spread remain scarce (6–8).

Mathematical modeling of HBV infection and treatment plays an important role in elucidating the interactions between HBV and its host (9–13). Recent modeling of HBV propagation from infection to steady state in humanized mice has emphasized the significance of viral production cycles within individual cells for understanding the complex, multiphasic kinetics of HBV infection *in vivo* (14). However, to date, there has been no modeling of HBV spread in PHH.

In this study, we aimed to characterize the kinetics of HBV infection by monitoring DNA at frequent time points and tracking HBV spread by monitoring the number of infected cells over 32 days post inoculation (p.i.) in PHH in the absence or presence of the HBV entry inhibitor Myr-preS1. Modeling was performed by adapting the recently developed ABM of HBV infection in humanized mice to simulate HBV-host interactions in PHHs (14), incorporating media replenishment and calibrating the model with measured HBV DNA and HBV-infected cell spread. We identified three distinct phases of both extracellular and intracellular HBV DNA kinetics. The ABM predicted that Myr-preS1 treatment is highly effective (91%) in blocking viral spread, as evidenced by an extremely slow increase in the third phase of both extracellular and intracellular HBV DNA kinetics. The model estimates also demonstrated that HBV infection dynamics in PHH in vitro closely mirrors that observed in the in vivo humanized liver chimeric mouse model.

## Results

### HBV infection of PHH *in vitro* reveals three extracellular HBV DNA kinetic phases

To characterize HBV kinetics in PHH up to 32 days p.i., 4 independent experiments were performed (see Materials and Methods). Extracellular HBV DNA was measured via qPCR revealing 3 kinetic phases consisting of a rapid decline (phase 1), a rapid increase (phase 2), and slow increase (phase 3) (**Fig. 1**, blue lines; **Table 1**). In Exp.1, there was a rapid decline phase with a slope of -0.82 log_10_/day (t_1/2_=8.8hr, 95% CI, [7.7, 10.2]) until day 3 p.i., followed by a rapid increase phase with a slope of ∼0.46 log_10_/day (t_2_=15.6hr, 95% CI, [14.7, 16.7]) between days 3-7 p.i., followed by a slower increase phase with a slope of ∼0.12 log_10_/day (t_2_=59.9hr, 95% CI, [53.3, 68.3]) between days 7-12 p.i. (**Fig. 1A**). Similar to Exp.1, the extracellular HBV DNA in Exp.2 increased rapidly with a slope of ∼0.51 log_10_/day (t_2_=14.2hr, 95% CI, [13.4, 15.0]) from day 3-7 p.i., followed by a slower increase with a slope of ∼0.15 log_10_/day (t_2_=49.6hr, 95% CI, [41.8, 61.0]) from days 7-12 p.i. (**Fig. 1B**). In Exp.3, a slow increase was seen from days 12-32 p.i. with a slope of ∼0.09 log_10_/day (t_2_=82.5hr, 95% CI, [78.6, 86.7]) (**Fig. 1C**). In Exp.4, which was initiated at lower MOI to provide a detailed longitudinal kinetic picture, the initial decline had a slope of -0.18 log_10_/day (t_1/2_=40.1hr, 95% CI, [35.0, 46.9]) until day 5 p.i.. This was followed by an increase with a slope of ∼0.17 log_10_/day (t_2_=42.8hr, 95% CI, [40.54, 45.29]) between days 5-17 p.i.. Thereafter, extracellular HBV DNA had a slower increase with a slope of ∼0.07 log_10_/day (t_2_=102.7hr, 95% CI, [93.3, 114.3]) from days 17-32 p.i. (**Fig. 1D**). Thus, as observed in our previous *in vivo* HBV infection in chimeric mice, the kinetics of extracellular HBV DNA was delayed in Exp.4 (GE=1) compared to the Exps.1-3 where the inoculum was a log higher (GE=10).

**Fig. 1.**
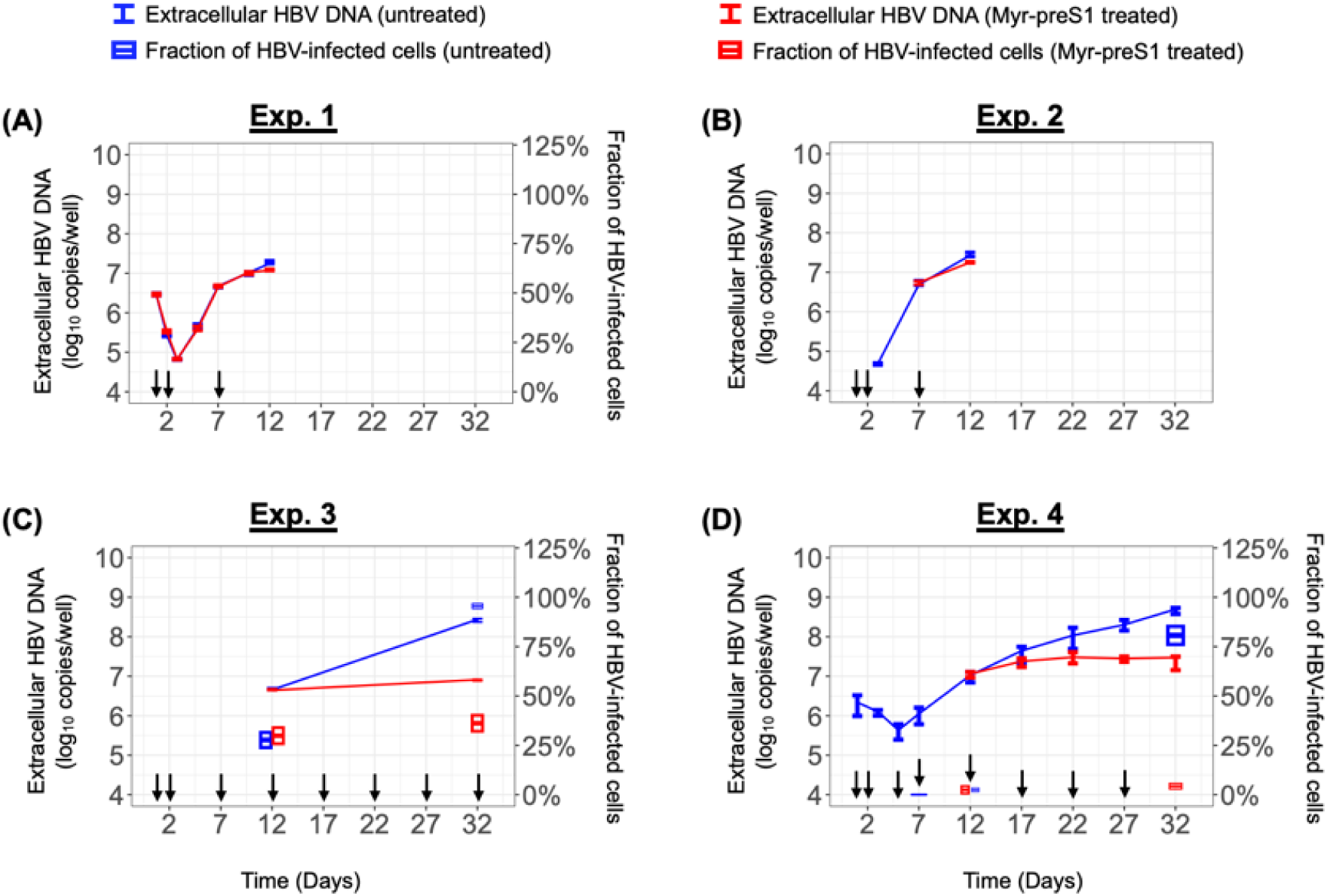
Measured excellular HBV DNA and percentage of HBV-infected cells in Exps. 1 – 4. **(A-D).** Solid line represents the median value of excellular HBV DNA in the control group (blue) and under Myr-preS1 treatment (red). Errorbar denotes the minimal and maximual measurments of excellular HBV DNA. Boxplot represents the percentage of HBV-infected cells (average±SD). Arrows indicate the media changes/replenishsment.

**TABLE 1.** Detailed characterization of extracellular HBV kinetics under Exps. 1-4.

| Exp.# | Phase 1<br>Slope (log<sub>10</sub>/d)<br>[95%CI] |  | Phase 2<br>Slope (log<sub>10</sub>/d)<br>[95%CI] |  | Phase 3<br>Slope (log<sub>10</sub>/d)<br>[95%CI] |  |
| --- | --- | --- | --- | --- | --- | --- |
|  | Untreated | Myr-preS1 | Untreated | Myr-preS1 | Untreated | Myr-preS1 |
| Exp.1 | -0.82<br>[-0.94, -0.71] | -0.93<br>[-1.00, -0.86] | 0.46<br>[0.43, 0.49] | 0.46<br>[0.42,0.50] | 0.12<br>[0.11, 0.14] | 0.09<br>[0.06, 0.11] |
| Exp.2 | NA | NA | 0.51<br>[0.48, 0.54] | NA | 0.15<br>[0.12, 0.17] | 0.10<br>[0.09, 0.12] |
| Exp.3 | NA | NA | NA | NA | 0.09<br>[0.08, 0.09] | 0.01<br>[0.01, 0.02] |
| Exp.4 | -0.18<br>[-0.21, -0.15] | NA | 0.17<br>[0.16, 0.18] | NA | 0.07<br>[0.06, 0.08] | 0.02<br>[0.01, 0.02] |
| NA, not available; CI, confidence interval. |  |  |  |  |  |  |

### Intracellular HBV DNA kinetics during infection of PHH *in vitro* mirrors the same three phases as extracellular HBV DNA

To characterize HBV intracellular kinetics in PHH up to 32 days post inoculation (p.i.), cell lysates were collected at each time point in the 4 experiments and total HBV DNA was measured via qPCR. Analogous to what was observed for extracellular HBV DNA, three phases of cell-associated HBV DNA kinetics were identified: rapid decline (phase 1), rapid increase (phase 2), and slow increase (phase 3) (**Fig. 2**, black lines and **Table 2**). In Exp.1, cell-associated HBV DNA declined with a slope of -0.23 log_10_/day (t_1/2_=31.7hr, 95% CI, [28.4, 35.8]) from days 1-3 p.i.. Thereafter, intracellular HBV DNA increased rapidly with a slope of ∼0.38 log_10_/day (t_2_=19.1hr, 95% CI, [17.2, 21.5]) from days 3-7 p.i., followed by a slower increase phase with a slope of ∼0.10 log_10_/day (t_2_=74.3hr, 95% CI, [63.6, 89.4]) from day 7 onwards (**Fig. 2A**). Likewise, in Exp.2, the intracellular HBV DNA rapidly increased with a slope of ∼0.42 log_10_/day (t_2_=17.2hr, 95% CI, [16.9, 17.4]) from days 3-7 p.i., followed by a slower increase with a slope of ∼0.12 log_10_/day (t_2_=59.4hr, 95% CI, [54.0, 66.0]) until day 12 p.i.. (**Fig. 2B**). Similar to the slower amplification phase of Exps. 1 and 2, the intracellular HBV DNA increased slowly with a slope of ∼0.08 log_10_/day (t_2_=92.1hr, 95% CI, [86.8, 98.2]) from days 12-2 p.i. in Exp.3 (**Fig. 2C**). Although initiated at a lower MOI of 1, we observed a similar kinetics of cellular HBV DNA in Exp.4: first phase decline slope of -0.18 log_10_/day (t_1/2_=39.4hr, 95% CI, [33.0, 49.1]) until day 3 p.i., followed by a fast increase with a slope of ∼0.19 log_10_/day (t_2_=39.0hr, 95% CI, [35.8, 42.8]) between days 3-7 p.i., followed by a slower increase with a slope of ∼ 0.08 log_10_/day (t_2_=87.3hr, 95% CI, [82.2, 93.0]) from days 7-32 p.i. (**Fig. 2D**).

**Fig. 2.**
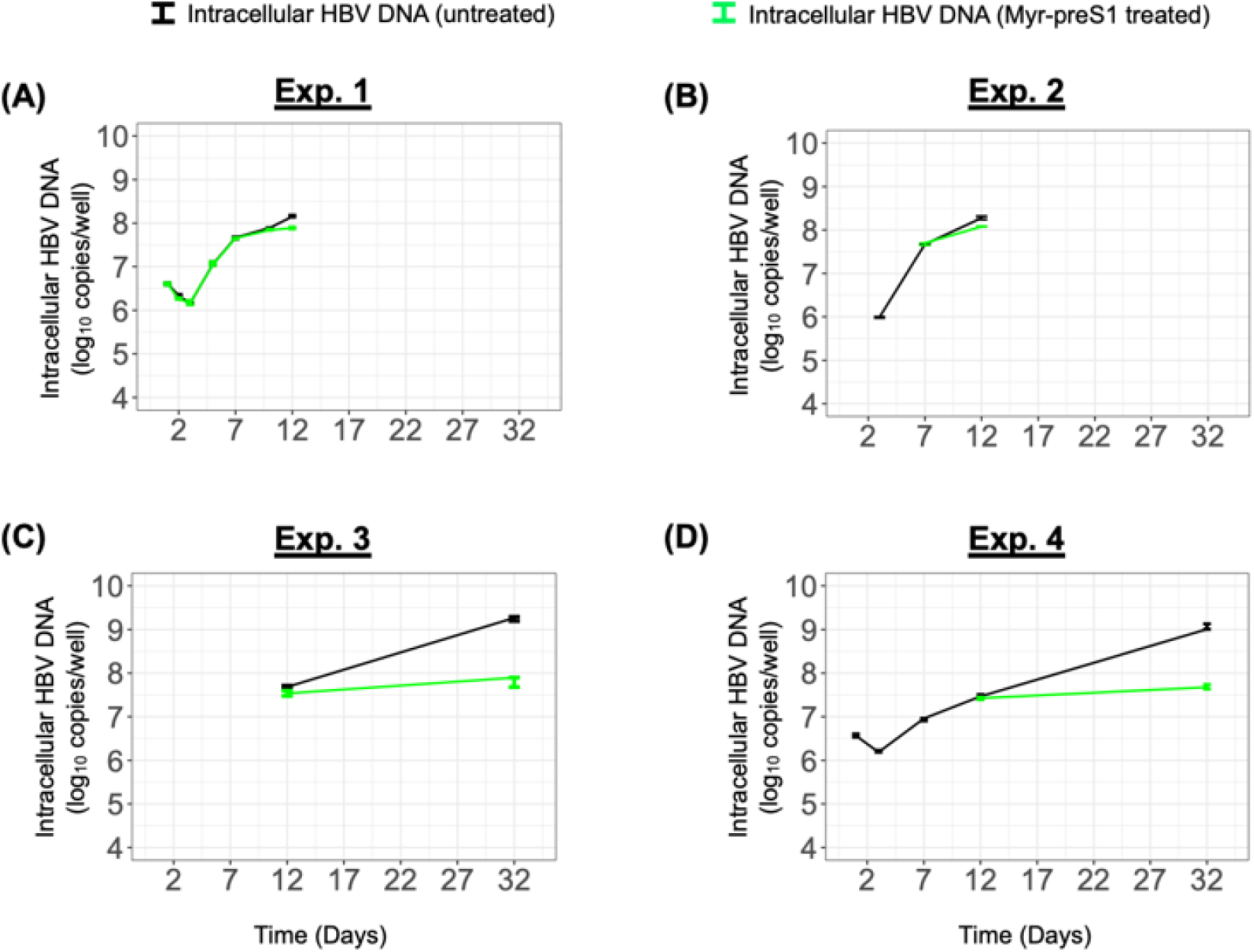
Measured intracellular HBV DNA in Exps. 1 – 4. **(A-D).** Solid line represents the median value of intracellular HBV DNA in the control group (black) and under Myr-preS1 treatment (green). Errorbar denotes the minimal and maximual measurments of intracellular HBV DNA.

**TABLE 2.** Detailed characterization of intracellular HBV kinetics under Exps. 1-4.

| Exp. # | Phase 1<br>Slope (log<sub>10</sub>/d)<br>[95%CI] |  | Phase 2<br>Slope (log<sub>10</sub>/d)<br>[95%CI] |  | Phase 3<br>Slope (log<sub>10</sub>/d)<br>[95%CI] |  |
| --- | --- | --- | --- | --- | --- | --- |
|  | Untreated | Myr-preS1 | Untreated | Myr-preS1 | Untreated | Myr-preS1 |
| Exp.1 | -0.23<br>[-0.25, -0.20] | -0.34<br>[-0.39, -30] | 0.38<br>[0.34, 0.42] | 0.37<br>[0.33, 0.41] | 0.10<br>[0.08, 0.11] | 0.05<br>[0.04, 0.06] |
| Exp.2 | NA | NA | 0.42<br>[0.41, 0.43] | NA | 0.12<br>[0.11, 0.13] | 0.08<br>[0.07, 0.09] |
| Exp.3 | NA | NA | NA | NA | 0.08<br>[0.07, 0.08] | 0.01<br>[0.00, 0.03] |
| Exp.4 | -0.18<br>[-0.22, -0.15] | NA | 0.19<br>[0.17, 0.20] | NA | 0.08<br>[0.08, 0.09] | 0.01<br>[0.01, 0.02] |
| NA, not available; CI, confidence interval. |  |  |  |  |  |  |

### Myr-preS1 treatment affects the third phase of both intracellular and extracellular HBV DNA kinetics

To determine the effect of blocking NTCP-mediated HBV entry in the 4 experiments performed, parallel cultures were treated with Myr-preS1 to block NTCP-mediated HBV entry starting 24h after HBV inoculation. Under Myr-preS1 treatment, extracellular HBV DNA kinetics in Exp.1 followed the same three phases: (phase 1) rapid decline with a slope of -0.93 log_10_/day (t_1/2_=7.8hr, 95% CI, [7.2, 8.4]) until day 3 p.i.; (phase 2) fast increase with a slope of ∼ 0.46 log_10_/day (t_2_=15.7hr, 95% CI, [14.4, 17.3]) between days 3-7 p.i.; and (phase 3) a slower increase with a slope of ∼0.09 log_10_/day (t_2_=84.8hr, 95% CI, [67.0, 115.6]) from day 7 onwards (**Fig. 1A**, red line). However, the last phase viral increase appeared slower and/or plateaued under Myr-preS1 treatment (**Fig. 1A**, red line), reduced from the ∼0.12 log_10_/day (t_2_=59.9hr, 95% CI, [53.3, 68.3]) increase observed in the absence of Myr-S1 (**Fig. 1A**, blue line). The same pattern was observed in Exp.2, with the slope between days 7-12 p.i. reduced from ∼0.15 log_10_/day (t_2_=49.6hr, 95% CI, [41.8, 61.0]) to 0.10 log_10_/day (t_2_=69.7hr, 95% CI, [59.8, 83.5]) in the presence of Mry-preS1 (**Fig. 1B**, blue vs. red line). Looking at days 12-32 p.i. in Exp.3, there was a much slower increase in extracellular HBV DNA in the presence Mry-preS1 (slope=0.01 log_10_/day; t_2_=583.6hr, 95% CI, [490.9, 719.5]) compared to untreated cultures (slope=0.09 log_10_/day; t_2_=82.5hr, 95% CI, [78.6, 86.7]) (**Fig. 1C**). Likewise, in Exp.4, the third phase amplification was remarkably slower in the presence of Myr-pres1 (slope=0.02 log_10_/day; t_2_=426.4hr, 95% CI, [324.7, 621.1]) compared to that observed in the untreated cultures (slope = 0.07 log_10_/day; t_2_=102.7hr, 95% CI, [93.3, 114.3]) (**Fig. 1D**).

To determine how blocking NTCP-mediated HBV entry in the 4 experiments impact intracellular HBV DNA kinetics, cell lysates were harvested from the Myr-preS1 treated cultures in all 4 experiments and cell-associated HBV DNA was measured via qPCR. Under Myr-preS1 treatment, intracellular HBV kinetics in Exp.1 follow three phases: (1) rapidly decline with a slope of -0.34 log_10_/day (t_1/2_=20.95hr, 95% CI, [18.54, 24.08]) until day 3 p.i.; (2) fast increase with a slope of ∼ 0.37 log_10_/day (t_2_=19.5hr, 95% CI, [17.6, 22.0]) between days 3-7 p.i.; (3) slow increase with a slope of ∼0.05 log_10_/day (t_2_=147.8hr, 95% CI, [119.2, 194.6]) from day 7 onwards. While Phase 1 and 2 were equivalent to the untreated cultures, Phase 3 had a considerably lower slope under Myr-preS1 treatments. The same pattern was observed in the other experiments. In Exp.2, intracellular HBV kinetics had a slower increase under Mry-preS1 treatment with a slope of 0.08 log_10_/day (t_2_=91.0hr, 95% CI, [79.4, 106.6]) between days 7-12 p.i.. In Exp.3, an extremely slow increase during phase 3 was observed under Myr-preS1 compared to untreated PHH (slope=∼0.01 log_10_/day; t_2_=501.20hr, 95% CI, [282.56, 2215.51] vs. ∼0.08 log_10_/day t_2_=92.10hr, 95% CI, [86.76, 98.14]) (**Fig. 1C**). In Exp.4, the Phase 3 slope under Mry-preS1 was 0.01 log_10_/day; t_2_=551.9hr, 95% CI, [407.8, 853.8] compared to the untreated culture Phase 3 slope ∼ 0.08 log_10_/day (t_2_=87.3hr, 95% CI, [82.2, 93.0]) (**Fig. 2D** and **Table 2**).

### Effect of Myr-preS1 reveals that the third phase of extracellular HBV infection in PHH *in vitro* represents viral spread

Because Myr-preS1 is expected to block HBV spread to new PHH, we fixed parallel cultures at the indicated time points in Exp.3 and Exp.4 to quantify the percentage of HBV-infected cells in untreated and Myr-preS1 treated cultures (**Fig. 1**, blue versus red boxplots, respectively). In Exp.3 on day 12 p.i., the mean percentage of HBV-infected cells in Myr-preS1-treated and untreated cultures was equivalent (29.9% ± 4.1% vs. 27.7% ± 3.9, respectively) (**Fig. 1C**). However, the percentage of HBV-positive cells was statistically lower under Myr-preS1 treatment compared to the untreated group on day 32 p.i. (36.2% ± 4.1% vs. 95.6% ± 0.4%, respectively) (**Fig. 1C**). Similarly, in Exp.4 there was no difference in the percentage HBV-infected cells between Mry-preS1-treated and untreated cells on day 12 p.i. (1.8% ± 1.9% vs. 2.0% ± 1.0%, respectively), but was a significant difference in the percentage of HBV-infected cells on day 32 p.i. (4.3% ± 1.4% vs. 80.7% ± 4.6%) (**Fig. 1D**). Together this demonstrates that Phase 3 primarily represents NTCP-dependent HBV spread.

### Model reproduces the kinetics of HBV infection observed in PHH and suggests high Myr-preS1 efficacy in blocking HBV spread

To capture the kinetics of HBV *in vitro* in PHH, we adapted our recently developed ABM and calibrated it with extracellular HBV data measured in Exp.4. Because Exp.4 has the most complete HBV infection kinetics within a single experiment, we assessed whether the model could fit the HBV kinetics observed in Exp.4, and the model reproduced well the extracellular HBV DNA multiphasic pattern (**Fig. 3A** and **3B**) observed as well as the percent infected cell data (**Fig. 3C** and **3D**). In addition, the oscillatory pattern of extracellular HBV DNA kinetics predicted in the ABM reflected well the media changes when the extracellular HBV DNA dramatically declined (**Fig. 3A** and **3B**). The ABM was calibrated, resulting in sampled joint posteriors (**Fig. S1**, upper and lower rectangles) and marginal parameter distributions (**Fig. S1**, diagonal rectangles) of all the model parameters, both non-intervention and Myr-preS1 intervention parameters. We used a calibration stopping criterion of 1000 effective sample size within the final calibration target bounds. These results reveal the parameter ranges for HBV infection and production and their high probability regions within those ranges that correspond to simulated target values consistent with the empirical targets. They also depict correlations between the infection-associated parameters (e.g., between ρ and β).

**Fig. 3.**
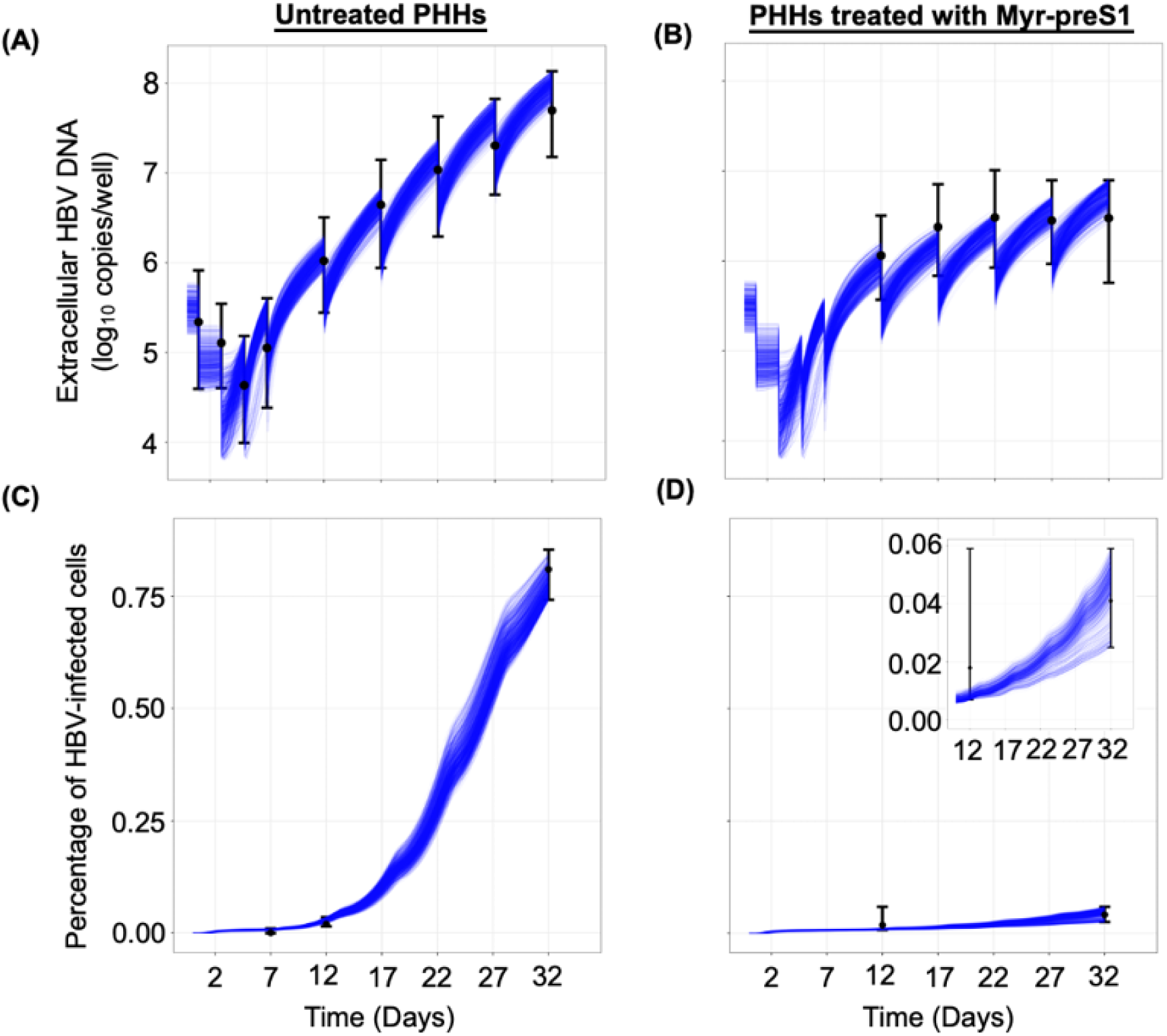
ABM calibration by fitting to Exp. 4. **(A-D).** ABM simultaneous calibrations (blue solid curves) with measured extracellular HBV DNA (top) and percentage of HBV-infected PHH (bottom) in Exp. 4. <u>Left panels (A & C)</u>: Untreated. <u>Right panels (B & D)</u>: Myr-preS1 treated. The ABM calibrations were done using the IMABC algorithm. Black dots are the median empirical calibration target values (i.e., extracellular HBV DNA) for each time point, error bars are the minimal and maximal bounds for the empirical targets, and blue lines represent 1000 IMABC posterior outputs, all of which are contained with all of the empirical calibration bounds.

These modeling results estimate that the viral intracellular eclipse phase lasts between 18-38 hours (**Table 3**). This non-productive eclipse phase ends and the P phase begins when the model predicts an increase in the intracellular viral production characterized by faster production cycles that release larger amounts of virus (**Fig. 4**). Specifically, during the first 2.5 days p.i., the model predicted that infected PHH have a long production cycle of 1 virion per day, but by 4 days p.i., infected PHH produce virions faster with a rate of 2 virion per hour. During the eclipse stage, virion production/release is minimal, so extracellular HBV DNA levels are observed to decrease consistent with the media removal/replenishment occurring at days 1 and 3 p.i.. After day 4 p.i., the virion production in each cell reaches a steady-state production rate of 4 virions per hour leading to an increase in virion release and infection of new PHH that explains the fast extracellular HBV DNA increase observed between days 5-17 p.i.. Then, as the number of target PHH decreases the extracellular HBV DNA increase slows consistent with the third phase of slower amplification. Again, looking at Exp.4, the model recapitulated the much slower increase in extracellular HBV DNA from days 12-32 p.i. under Myr-preS1 treatment (**Fig. 3B** and **3D**). Based on the calibration, the ABM predicts a median Myr-preS1 efficacy of 91.0% (first quantile [Q1]: 90% - third quantile [Q3]: 92%) to be achieved after a median 54 hours (Q1:40 hr – Q3:65 hr) post treatment initiation, suggesting a high efficacy of Myr-preS1 treatment in blocking HBV spread at an early stage of infection.

**Fig. 4.**
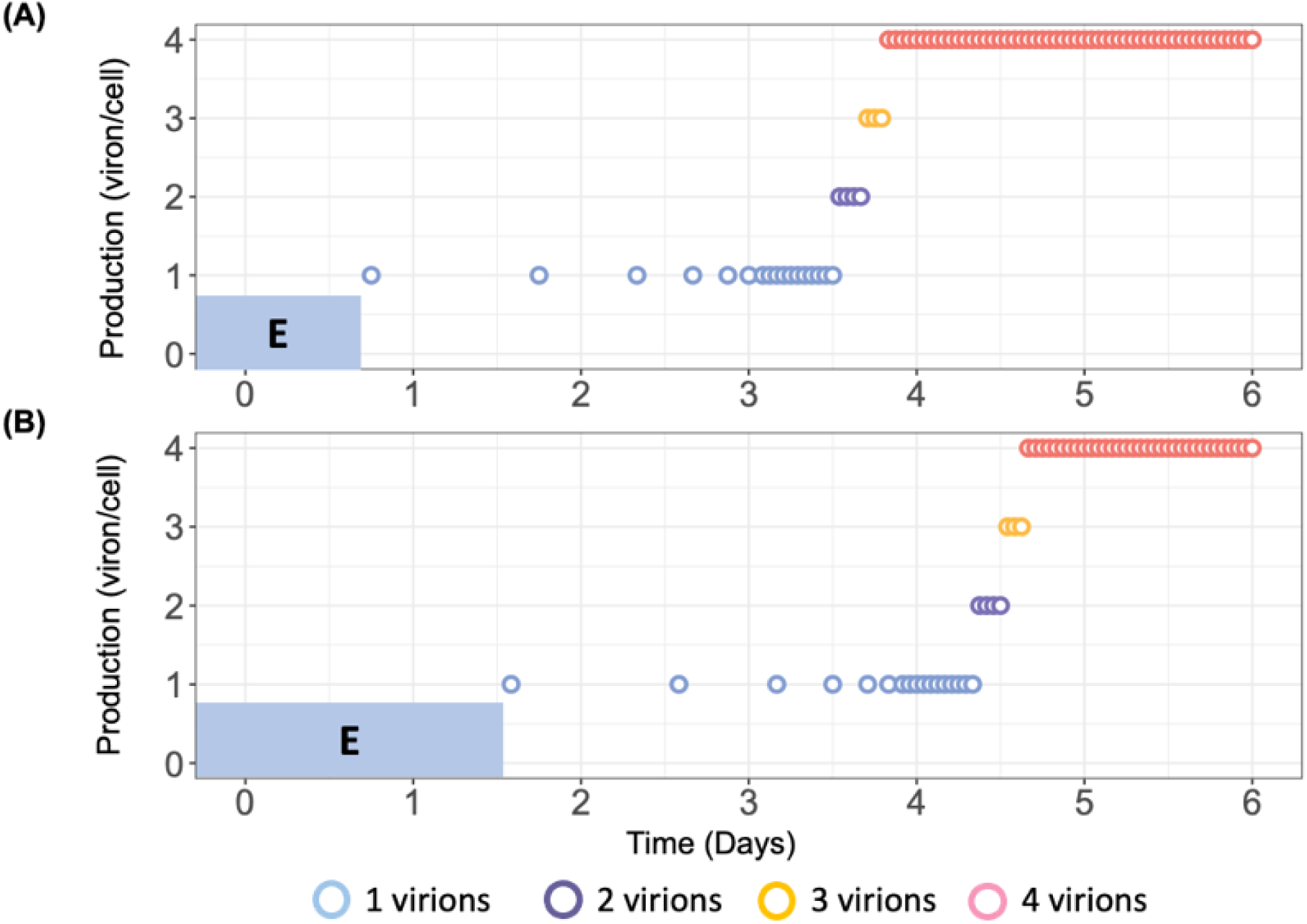
ABM prediction of HBV virion production cycles in PHHs. Blue-shaded “E” represents minimal eclipse phase **(A)** and maximal eclipse phase **(B)** for PHHs. Each PHH has a randomized eclipse phase following by a consistent virion production pattern starts with 1 virion/day. As virus resources accumulate, the production cycle shortens. Production also increases to 2, and then 3 virions before reaching steady state of 4 virions/hr. The magnitude of virion production was calculated using **Eq. 1** and the time between each production cycle was calculated using **Eq. 2**. The parameter values (α, P_st_, ψ, δ, ω) used to calibrate **Eq. 1** and **Eq. 2** were the median level of estimated parameters (**Table 4**).

**TABLE 3.** IMABC calibration for fitting Exp. 4 with and without Myr-preS1. Each parameter was sampled based on a uniform distribution (prior distribution) to implement IMABC. IQR represents the difference between the 25^th^ and 75^th^ percentiles of the marginal posterior distribution of parameters. All IMABC fits are shown in Fig. 3.

| Category | Parameter description | Symbol [unit] | Prior simulations and preliminary fits (uniform) |  | IMABC calibration |  |
| --- | --- | --- | --- | --- | --- | --- |
|  |  |  | [min-max] | Source | median | IQR [Q1-Q3] |
| **<u>Initial condition</u>** | Initial uninfected PHH number | $T_0$ [cell] | [38000-42000] | Expert opinion<sup>1</sup> | 39900 | [38874-41040] |
| | Initial viral load | $V_0$ [virion] | [15924-100475] | Expert opinion<sup>1</sup> | 30662 | [24779-37093] |
| **<u>Infection</u>** | Fraction of virion that are infectious | $\rho$ | [0.145-0.58] | (14) | 0.28 | [0.19-0.33] |
| | Infection rate | $\beta$ [cell/hr] | [0.0001-0.0025] | (14) | 0.0009 | [0.0005-0.0011] |
| **<u>Eclipse phase</u>** | Min eclipse phase | $\Omega_{min}$ [hr] | [5-36] | (14) | 18 | [12-26] |
| | Max eclipse phase | $\Omega_{max}$ [hr] | [18-50] | (14) | 38 | [30-45] |
| **<u>Production (Eq. 1 and Eq.2)</u>** | Initial production cycle length | $\delta$ [hr] | [15-30] | (14) | 23.41 | [19.62-27.19] |
| | Decay constant | $\omega$ [hr<sup>-1</sup>] | [0.3-0.97] | (14) | 0.54 | [0.50-0.60] |
| | Number of cycles to reach 50% of $P_{st}$ | $\alpha$ [hr] | [10-30] | (14) | 21 | [15-26] |
| | Steepness of the production curve | $\gamma$ [hr<sup>-1</sup>] | [0.1-0.9] | (14) | 0.35 | [0.26-0.48] |
| | Virus production at steady state | $P_{st}$ [virion/cell/hr] | [2-5] | (14) | 4 | [3-4] |
| **<u>Media change</u>** | Removal rate | $R_r$ | [0.01-1] | Educational fits<sup>2</sup> | 0.75 | [0.68-0.77] |
| **<u>Myr-preS1 treatment</u>** | Efficacy of Myr-preS1 | $\eta$ | [0.5-1] | Educational fits<sup>2</sup> | 0.91 | [0.90-0.92] |
| | Time at Myr-preS1 taking effect | $t_{eff}$ [hr] | [24-72] | Educational fits<sup>2</sup> | 54 | [40-65] |

Note:
1. Our experimentalists suggest that initial uninfected PHH number ( $T_0$ ) in an individual well is from $3.8 \times 10^5$ to $4.2 \times 10^5$ . To improve the computational efficiency, we used 1/10 of the actual number which is the range of $3.8 \times 10^4$ to $4.2 \times 10^4$ in the ABM to model the virus spread in the culture media. The initial viral load ( $V_0$ ) is $4.0 \times 10^5$ per well which was rescaled to $4.0 \times 10^4$ to conform to the scaled-down PHH number. In addition, 0.4log was added to the lower and upper bounds of each viral measurement to account for experimental variations so the final range for the initial viral load is from 15924 (4.2log) to 100475 (5log). 2. The range was determined based on preliminary model fittings using AnyLogic optimization tool.

**TABLE 4.** Initial conditions for each experiment.

|  | Exp.1 | Exp. 2 | Exp.3 | Exp.4 |
| --- | --- | --- | --- | --- |
| Number of Hepatocyte<br>(cells/well) | 2.0 x 10<sup>5</sup> | 4.0 x 10<sup>5</sup> | 1.6 x 10<sup>5</sup> | 4.0 x 10<sup>5</sup> |
| Plate<br>(well) | 24 | 24 | 48 | 24 |
| Volume of culture medium<br>(uL/well) | 500 | 500 | 200 | 500 |
| MOI<br>(genome equivalents/cell) | 10 | 10 | 10 | 1 |

## Discussion

In this study, we characterized the viral kinetics of HBV infection and assessed viral spread over a period of 32 days in PHH. We identified three distinct phases in both extracellular and intracellular HBV DNA kinetics: a first rapid decline, a second rapid increase, and a third slow increase phase. Notably, the second rapid and third slow increase phases observed in PHH closely resemble the two amplification phases (phase 4 and 6) seen in the humanized mouse model (14). We show that the ABM reproduces well the measured viral DNA kinetic and spread data by assuming a cyclic nature of viral production, characterized by an initial slow but increasing rate of HBV production in individual cells, consistent with the modeling findings during acute HBV infection in humanized mice (14). The ABM also predicts that Myr-preS1 treatment is highly effective (91±1%) in blocking viral spread, as evidenced by an extremely slow increase in the third phase of both extracellular and intracellular HBV DNA kinetics.

Although extracellular and intracellular HBV DNA exhibited similar kinetic patterns across 4 experiments, a more detailed analysis of viral kinetics under varying HBV inocula dose revealed notable differences. First, the extracellular HBV DNA level in Exp.1 declined approximately five times faster (−0.82 log10/day) than in Exp.4 (−0.18 log10/day) (**Table 1**). Given that that HBV degradation is relatively stable in media (15), the faster decline in Exp.1 might suggest an unexpectedly higher rate of HBV removal from the media by media replenishment. In addition, the rapid increase phase in Exp.1 occurred ∼3 times faster (0.46 log_10_/day) compared to Exp.4 (0.17 log_10_/day) (**Table 1**), Because higher inoculum HBV levels (GE=10 in Exp.1 > GE=1 in Exp.4) is expected to result in higher viral uptake, this suggests that a higher initial HBV levels (perhaps within individual cells) may lead to more efficient virion production and/or faster viral spread, as evidenced by a two-fold faster rapid increase phase of intracellular HBV DNA in Exp.1 (0.38 log_10_/day) compared to Exp.4 (0.19 log_10_/day) (**Table 2**).

To the best of our knowledge, HBV spread in PHH has not been investigated in previous studies. The lack of increase in extracellular HBV DNA, intracellular HBV DNA, and the number of infected cells during phase 3 in the presence of Myr-preS1 suggests that phase 2 reflects viral amplification within initially infected cells, while phase 3 represents the spread to naïve cells. Consequently, phase 3 of intracellular HBV amplification, in the absence of Myr-preS1, parallels the spread observed in extracellular HBV kinetics. The continued, though slower, increase in intracellular HBV levels (with a slope of 0.01-0.08 log10/day; Table 2) during phase 3 in the presence of Myr-preS1 suggests either incomplete inhibition of viral spread or that intracellular HBV levels had not yet reached steady state.

To better understand the HBV kinetics observed in PHH, we adapted our recently developed ABM by incorporating media replenishment and accounting for the impact of Myr-preS1 interventions on HBV spread. We estimated several key parameters that reflect both the duration of the HBV eclipse phase and its subsequent production cycles. Interestingly, our results showed that the duration of the HBV eclipse phase in PHH ranges from 18 to 38 hours, a more narrow and consistent range compared to the 7-50 hour duration predicted for the chimeric mouse model (14). The model’s estimate for virion production at steady state (i.e., P_st_) is identical in both systems, with approximately 4 virions per infected cell per hour in PHH and in mice. Other key parameters related to the viral production cycle, such as initial production cycle length (20-27hr in PHH vs. 23-30hr in humanized mice), the number of viral production cycles to reach 50% of P_st_ (15-26 in PHH vs. 14-30 in humanized mice), and the steepness of the production curves (0.50-0.60hr^-1^ in PHH vs. 0.40-0.60hr^-1^ in humanized mice), also show similarities, further suggesting that HBV infection kinetics in PHH closely resemble those in the chimeric mouse model. This similarity indicates that viral replication is similarly regulated in human hepatocytes *in vitro* and *in vivo*. While it is acknowledged that *in vitro* experiments lack the physiological benefits of *in vivo* systems—such as the influence of proteins from other organs, including cytokines, chemokines, and hormones—these *in vitro* models offer a valuable platform for analyzing the detailed dynamics of HBV replication, free from the confounding effects of such factors.

HBV infects hepatocytes by first attaching to heparan sulfate proteoglycans (HSPGs) and subsequently binding to the HBV receptor, human sodium taurocholate co-transporting polypeptide (hNTCP). We employed treatment with Myr-preS1 entry inhibitor, to block viral binding to hNTCP by competitive inhibition. To assess its efficacy in PHH, Myr-preS1 was administered starting one day post-inoculation to prevent the spread of HBV infection. The model predicts that the fraction of HBV-infected cells increased very slowly, from approximately 2-7%, between days 17-32 p.i., suggesting that Myr-preS1 effectively interfered with HBV spread (**Fig. 3D**). The estimated median efficacy of Myr-preS1 treatment was 91%, indicating that (1) higher dosages may be required to fully block HBV infection, (2) HBV may interact with other receptors during cell entry, and/or (3) additional spread mechanisms (e.g., spread to adjacent cells, forming infected cell clusters (16)), may contribute to further HBV infection at later stages. The lack of decline in intracellular HBV DNA under Myr-preS1 treatment suggests that HBV infection remains stable in the absence of re-infection, which also implies that cccDNA maintenance is not dependent on re-infection under these conditions. However, with the cccDNA half-life predicted to be approximately 40 days in vitro, longer-term experiments are needed to confirm these findings.

Our study has several limitations. First, we used only Exp.4 for modeling, as Experiments 1 to 3 had less frequent data sampling, which limited our ability to model the differences in HBV kinetics at varying HBV input doses. Future experimental work with more frequent sampling points is needed to compare HBV viral kinetics across different doses. Second, we did not include intracellular HBV DNA or cccDNA in the model for the sake of simplicity. To gain a deeper understanding of intracellular HBV dynamics, an integration of the current ABM with our recent mathematical model that accounts for intracellular viral DNA and cccDNA kinetics (15) is warranted. Lastly, PHH death was not incorporated into the ABM. In our PHH culture system, although cell density slightly decreased by day 2 p.i., the number of hepatocytes remained stable for at least 33 days (17). Based on these observations and for model simplification, we assumed that excluding PHH loss or death would not significantly affect the results. However, for longer-term experiments monitoring HBV infection in PHH, the inclusion of cell death and PHH loss may be necessary for accurately calibrating HBV kinetics in the model.

In conclusion, this study demonstrates that HBV DNA levels follow multiphasic replication kinetics in PHH. Myr-preS1 treatment effectively blocks HBV spread, although it does not prevent it completely. By comparing model predictions from both the in vivo humanized mouse model and the in vitro PHH system, we show that HBV infection dynamics in PHH closely mirror those observed in the uPA/SCID chimeric mouse model, suggesting that there are virtually no other factors in this mouse model that impacting HBV spread. Future studies, including mathematical modeling of intracellular replication and cccDNA transcription, will provide deeper insights into HBV spread and HBV-host dynamics at the molecular level.

## Materials and Methods

### PHH Preparation

Commercially available cryopreserved human hepatocytes (BD Bioscience, lot195, Hispanic, female, 2 years) were transplanted into cDNA-uPA/SCID mice via injection into the spleen. PHH were isolated from the chimeric mice with humanized livers at 9-15 weeks after transplantation by a standard collagenase perfusion. To expand the cells, the isolated PHH were serially transplanted into additional cDNA-uPA/SCID mice as previously described (18). Nine to fifteen weeks after the serial transplantation, PHH were collected for in vitro culture and infection. PHH were seed in collagen-coated 24 well plates (Corning, Japan) at 2.0×10^5^ cells/well (Exp.1) and 4.0×10^5^ cells/well (Exp.2 and 4) or in 48 well plates at 1.6×10^5^ cells/well (Exp.3) and incubated at 37C° with 5% CO_2_ in dHCGM medium (**Table 4**). The base of this medium is Dulbecco’s modified Eagle’ medium (Gibco ThermoFisher Scientific) supplemented with 44mM NaHCO3 (Wako Chemicals), 100IU/ml Penicillin G (Invitrogen ThermoFisher Scientific), 100μg/mL Streptomycin (Invitrogen ThermoFisher Scientific), 20mM HEPES (Gibco ThermoFisher Scientific), 10% FBS (Biosera), 15μg/mL L-proline (Wako Chemicals), 0.25μg/mL insulin (Sigma-Aldrich), 50nM Dexamethasone (Sigma-Aldrich), 5ng/mL Epidermal growth factor (Sigma-Aldrich), 0.1mM L-ascorbic acid 2-phosphate (Wako Chemicals), and 2% Dimethyl sulfoxide (Sigma-Aldrich).

### HBV Inoculum

HBV genotype C2 obtained from a chronic hepatitis B patient and named Hiroshima GtC CL3 (NCBI accession No. MH818373) was used as inoculum in this study. The virus was amplified in cDNA-uPA/SCID mice as previously described (18).

### HBV Infection

PHH were treated with an inoculum of 10 HBV GEq/cell (Exp.1-3) or 1 HBV GEq/cell (Exp.4) for 1 day with 4% polyethylene glycol 8000 (PEG 8000). The inoculated cells were cultured with 200 μL or 500 μL of culture medium for 48 well and 24 well plates, respectively. Myr-preS1 (6.25 μg/mL) treatment was initiated one day after inoculation and then continued throughout the experiment. Extracellular HBV DNA and intracelluar HBV DNA were measured in replicate wells at the indicated time points for each experiment. During the infection, culture media was renewed after HBV inoculation or until the end of the experiment as indicated by arrows in **Fig. 5**.

**Fig. 5.**
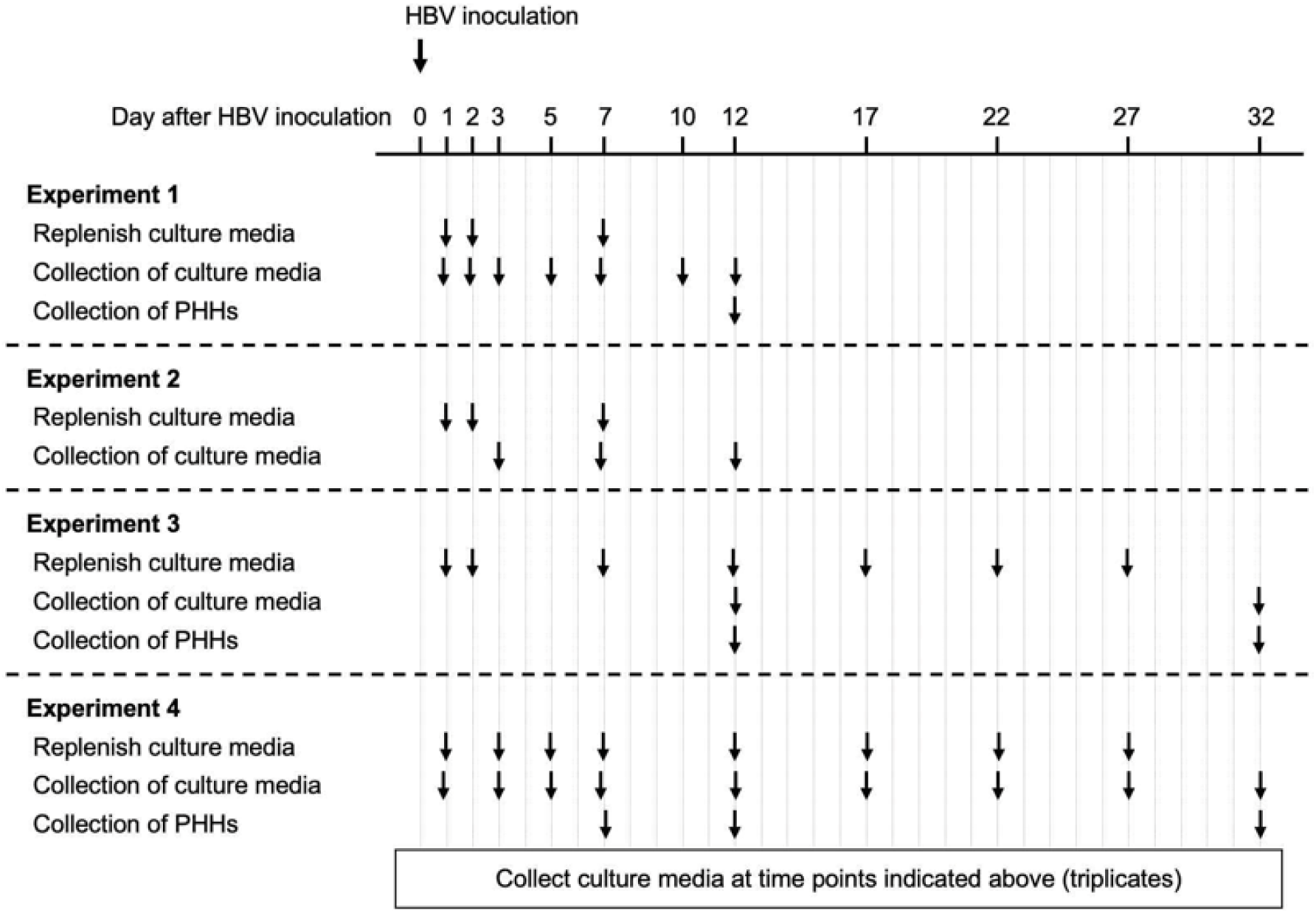
Experimental design for PHHs. PHHs were treated with an inoculum of 10 HBV GEq/cell (Exp 1-3) or 1 HBV GEq/cell (Exp 4) for 1 day, starting on day 0. Myr-preS1 (6.25 μg/mL) treatment was initiated one day after inoculation and then continued throughout the experiment. Extracellular HBV DNA and intracelluar HBV DNA were measured in collected culture media at the indicated time points for each experiment. PHHs were harvested for estimating HBV infected cells. During the infection, culture media was renewed after HBV inoculation or until the end of the experiment as indicated by arrows.

### Quantification of extracellular HBV DNA in culture medium

HBV DNA levels in culture media were quantified by qPCR as previously described (19). DNA was extracted using SMI TEST (Genome Science Laboratories, Tokyo, Japan) and dissolved in 20μL of H_2_O. Real-time PCR analysis was performed using ABI Prism 7300 Sequence Detection System. Amplification was performed in a 25μL reaction mixture containing SYBR Green PCR Master Mix (Applied Biosystems, Foster City, CA), 200nM of forward primer, 200nM of reverse primer, and 1μL of DNA or cDNA solution. After incubation for 2 minutes at 50°C, the sample was heated for 10 minutes at 95°C for denaturing, followed by a PCR cycling program consisting of 40 2-step cycles of 15 seconds at 95°C and 60 seconds at 60°C. The lower detection limit of this assay was 2.3 log copies/ml. The primers used for extraction are as follows: forward 5’-TTTGGGGCATGGACATTGAC-3’ and reverse 5’-GAGTGCTGTATGGTGAGGTG-3’.

### HBV DNA isolation and analysis

DNA was extracted from the harvested PHH using SMITEST (Genome Science Laboratories, Tokyo, Japan) in accordance with the manufacturer’s instructions. Intracellular HBV DNA and cccDNA levels were quantified by qPCR and TaqMan PCR, respectively, using an ABI Prism 7300 (Applied Biosystems, Carlsbad, CA, USA), as previously described (20). Briefly, the concentration of purified DNA was measured by BioPhotometer® 6131 (Eppendorf, Tokyo, Japan). Intracellular HBV DNA was quantified by qPCR using the same protocol and primers for extracellular HBV DNA quantification. Single-stranded DNA was selectively degraded from the extracted DNA using S1 nuclease (TakaraBio, Shiga, Japan), and then cccDNA levels were quantified. For total HBV DNA the forward primer (nucleotides [nt] 1521-1545: 5-GGGGCGCACCTCTCTTTACGCGGTC-3), reverse primer (nt 1862-1886: 5-CAAGGCACAGCTTGGAGGCTTGAAC-3), and TaqMan MGB probe (nt 1685-1704: 5-FAM-AACGACCGACCTTGAGGCAT-MGB-3) were utilized. PCR was performed using 100ng of extracted DNA, TaqMan® Fast Advanced Master Mix (Applied Biosystems), 300nmol of each primer and 250nmol of the probe. Amplification was performed as follows: 50°C for 2 min, then 95°C for 10 min, followed by 45 cycles of 95°C for 10 sec, 58°C for 5 sec, 63°C for 10 sec, and 20 sec at 72°C. The lower limited detection was 100 copies/100ng DNA.

### Estimation of HBV infected cells

Immunostaining for HBsAg was conducted as previously described (18). At indicated times after inoculation, PHH were fixed with 10% formalin neutral buffer solution for 10 minutes at room temperature. After washing twice with PBS, the fixed PHHs were treated with anti-HBsAg goat polyclonal antibody (bs-1557G; Bioss Inc, Woburn, MA). To analyze the percentage of HBsAg-positive PHH, five pictures were taken with a BZ-X700 microscope (Keyence, Osaka, Japan) and the number of PHH and HBsAg-positive PHH were counted.

### Modeling HBV dynamics

To study HBV spread in PHH, we modified our recently developed ABM to describe HBV infection in humanized mice (14). Analogous to the published model, this ABM consists of two types of agents to account for the PHH and the extracellular virus. Here, the PHH are represented by a two-dimensional grid of stationary cells while the virus is represented by freely diffusing agents intended to represent the amount of HBV in the culture medium. Each individual PHH can be in one of the following three discrete states: uninfected/susceptible PHH targets (T), infected PHH in an eclipse phase (I_E_), or productively infected PHH secreting progeny virus (I_P_) (**Fig. 6**). The ABM execution is again an iterative process where each iteration represents a discrete time step, each step = 1 hour. For each iteration, uninfected/susceptible PHH are infected and initially enter an eclipse phase (I_E_). After a random period of time following a uniform distribution of *U*(Ω_min_, Ω_max_), the infected PHH proceeded from the eclipse phase to the productive phase, and then they start to release newly generated progeny virus (i.e., I_p_). The magnitude and frequency of secreted virus is calculated using the following two equations to mimic the cyclic nature of the viral lifecycle within each cell (**Eq.1** and **Eq.2**)

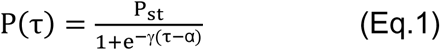

**Fig. 6.**
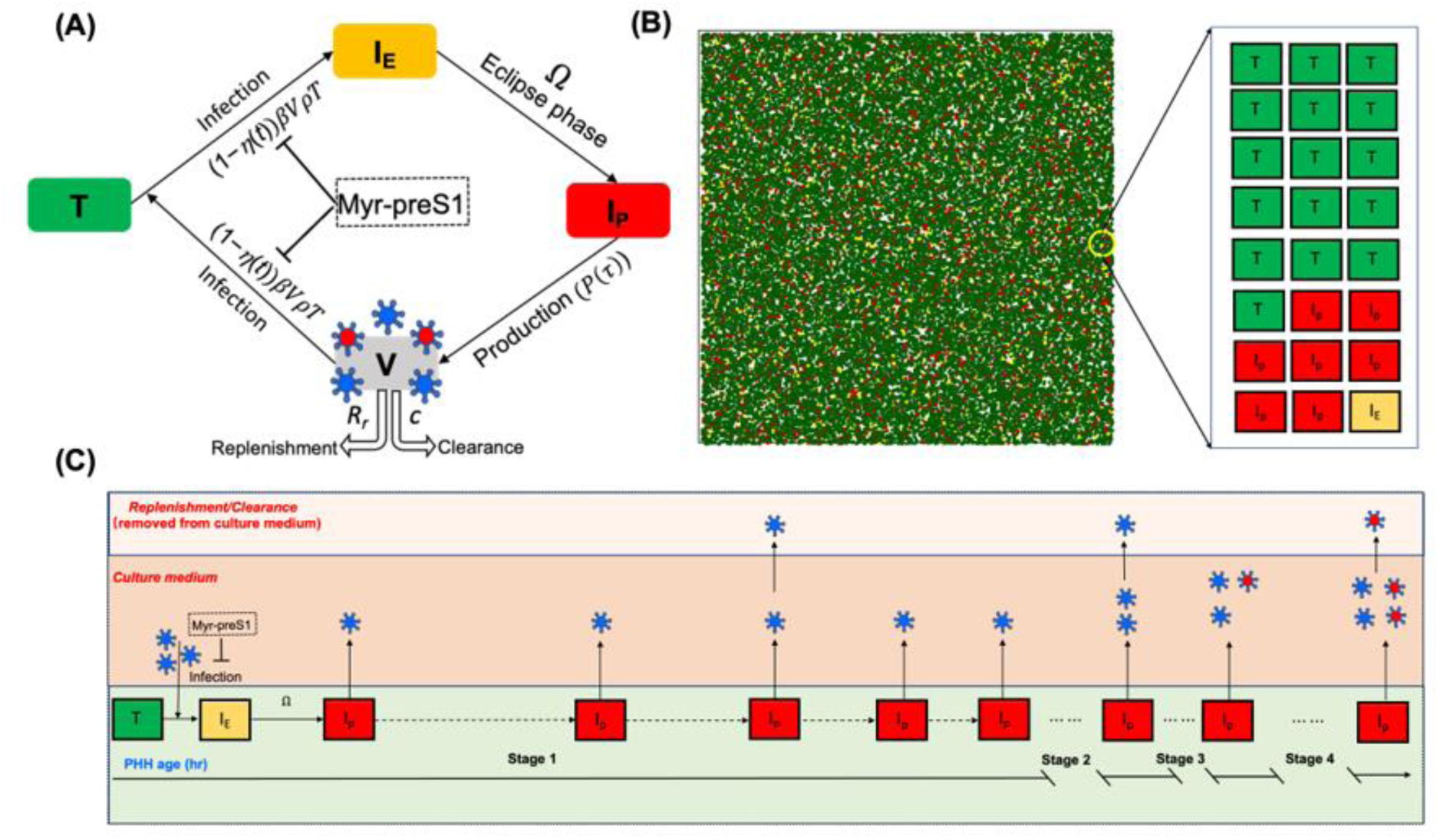
Conceptualiztion of the agent-based model. **(A)** The human hepatocytes can be only in one of the following three phases at a given time; T = uninfected cells which are termed as target or susceptible cells, I_E_ = HBV-Infected cells in eclipse phase (i.e., not yet releasing virions), I_P_ = HBV-infected cells actively producing/releasing virions. Once I_E_ become I_p_, they produce free virus that can further infect T. The free virus in blood, V, is composed of infectious (red center virus) and non-infections virions (blue center virus). The parameter ρ represents the fraction of virions that are infectious, *β* represents the infection rate constant, Ω represents eclipse phase duration, *P*(*τ*) represents virion secretion from I_P_ (**Eq. 1**), *c* represents viral clearance from blood, *R_r_* represents the portion of virions removed during medium change/replenishment, and we assume no death/loss for PHHs in the culture media. The effectiveness of Myr-preS1 is η when the drug takes effect at t_eff_. **(B)** Simulated hepatocytes including T (green), I_E_ (yellow), and I_P_ (red) in ABM at 15 days after HBV inoculation. **(C)** Schematic diagram of a representative hepatocyte progressing through ABM. Each individual hepatocyte has its own infection kinetics followed by a randomized eclipse phase and viral production cycle (Ω + Stage 1 + Stage 2 + Stage 3 + Stage 4). The virions were initially released by I_P_ starting with a long production cycle of 1 virion per cell (Stage 1: ∼0-2.5 days) that gradually reaches a production of 2 virions per cell with a shorten production cycle (Stage 2: ∼2.5-3 days) and then proceeds to 3 virions per cell (Stage 3: ∼3-4 days) before virion production increases to reach to a steady state production rate of 4 virions per hour per cell (Stage 4: ∼ 4 days onwards).

where *P*(*τ*) is number of virions produced by infected cells a *τ*, P_st_ is steady state virus production, α is number of cycles to reach to 50% of P_st_ , γ is steepness of the production curve, and *τ* is the production cycle

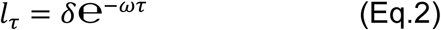

where l_τ_ is interval between production cycle, *τ* is the production cycle, *δ* is scaling factor indicating the initial production cycle length, and *ω* is decay constant.

To account for the removal of virus containing media and replenishment with fresh media, a removal/replenishment rate (*Rr*) was newly incorporated into the *in vitro* ABM which estimates the percentage of extracellular virus removed at each media change. To estimate the effect of the drug intervention on HBV spread, the following step function (**Eq.3**) was also incorporated into the ABM

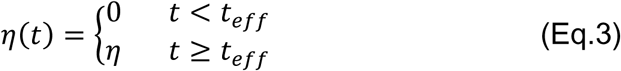

where η is efficacy of Myr-preS1 and t_eff_ is time when Myr-preS1 starts to take effect. A schematic picture of the modified agent-based model is shown in **Fig. 6**.

### Parameter estimation

We previously showed that the degradation of extracellular HBV DNA *in vitro* is extremely slow since the HBV is quite stable in the culture medium (21). Thus, in the ABM we assumed a fixed virus clearance rate of 0.0004 h^-1^ (t_1/2_ =72.2 days) in between the media changes. To estimate the remaining parameters, we applied the following two steps including preliminary AnyLogic fits and Incremental Mixture Approximate Bayesian Computation (IMABC) (22). Removal/replenishment parameter *Rr* was estimated to be in the range of (0.01-1) by fitting the ABM to the untreated PHH experimental data using AnyLogic optimization tool. Parameter ranges for efficacy of Myr-preS1 (η) and time when Myr-preS1 takes effect (t_eff_) were estimated by fitting the ABM to the experimental data under Myr-preS1 intervention. Parameter ranges for other parameters were determined based on the *in vivo* mouse model (14). By assuming uniform prior distributions for each parameter using the estimated ranges in the first step, we applied IMABC to obtain the posterior distributions of parameters.

### Model calibration using IMABC

The agent-based model (ABM) was calibrated by identifying parameters that result in model predictions that fit the experimental data. We utilized the IMABC algorithm (22) for calibration, implemented with the R IMABC package (23) and the EMEWS framework (24) on Argonne’s LCRC Bebop high-performance computing cluster. The IMABC algorithm works by adaptively constraining model output target bounds. The algorithm converges when a sufficient number of model parameters (effective sample size, i.e., 1000 samples) are found to simulate targets within the final empirical target bounds. During model calibration, two sets of ABM targets were sequentially evaluated, corresponding to non-intervention and intervention data. The calibration algorithm first used the ABM to simulate non-intervention targets (i.e., untreated PHH). If the non-intervention ABM did not result in points within the non-intervention targets, the parameter set was rejected. For each parameter vector that produced simulated targets (**Table S1**) within the specified IMABC target bounds, the algorithm next evaluated fit to intervention targets from the Myr-preS1 intervention. Through this approach the non-intervention and intervention ABM parameters were simultaneously estimated.

## Acknowledgements

This work was performed at the Natural Science Center for Basic Research and Development, Hiroshima University. This study was partly supported by Fund for the Promotion of Joint International Research (Fostering Joint International Research) from Japan Society for the Promotion of Science (Grant number: 17KK0194), the Japan Agency for Medical Research and Development (AMED; Grant numbers: JP20fk0310107s0504, fk0310513h0001 and JP20fk0310109h0004), the Japanese Ministry of Education, Culture, Sports, Science, and Technology (MEXT; Grant numbers: 18K07973 and 21k08006), and NIH (Grant numbers: R01AI144112 and R01AI146917). This research was completed with resources provided by the Laboratory Computing Resource Center at Argonne National Laboratory. The authors declare no conflict of interest pertaining to this study.

## Supplemental Material

**Table S1.** Experimental data (simulated targets) for Exp. 4.

| Day | Extra HBV-median [min, median, max] | Percentage of HBV infected cells [min, median, max] |
| --- | --- | --- |
| 1 | [4.595, 5.339, 5.915] |  |
| 3 | [4.603, 5.106, 5.542] |  |
| 5 | [3.994, 4.634, 5.184] |  |
| 7 | [4.384, 5.050, 5.604] | [0, 0.0005, 0.01] |
| 12 | [5.443, 6.021, 6.504] | [0.015, 0.02, 0.035] |
| 17 | [5.941, 6.644, 7.146] |  |
| 22 | [6.291, 7.034, 7.627] |  |
| 27 | [6.758, 7.304, 7.823] |  |
| 32 | [7.178, 7.696, 8.131] | [0.742, 0.810, 0.854] |

**Fig. S1.**
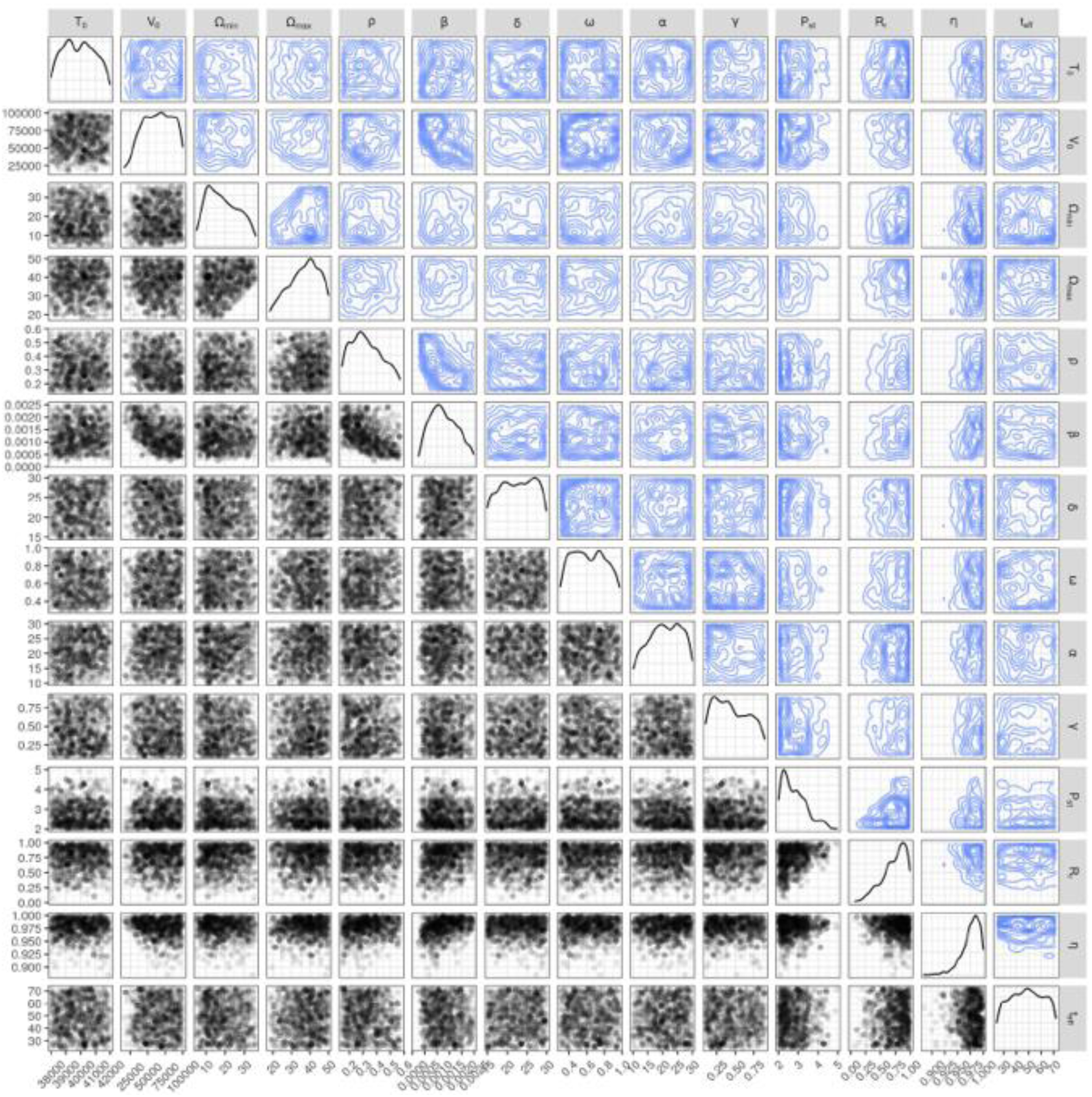
Joint posterior distributions of non-intervention and Myr-preS1 intervention parameters from IMABC (1000 samples). The upper (contour plots) and lower (scatter plots) triangles show the two-dimensional joint posterior distributions. The diagonal shows the marginal distributions of each parameter in **Table 3**.

